# Airborne DNA reveals synchronized responses of tropical forest assemblages to precipitation

**DOI:** 10.64898/2026.09.18.752615

**Authors:** Joseph M. Craine, Nicholas Schulte, Jessica Devitt, Devin Leopold, Mitchell Ralson, Kristin Saltonstall, Noah Fierer

**Author notes:** Corresponding author: Joseph Craine.

## Abstract

Tropical forests contain much of Earth’s biodiversity, yet community-wide responses to seasonal climate transitions remain poorly resolved. We used weekly airborne environmental DNA sampling, supplemented by spatially intensive dry- and wet-season campaigns, to characterize fungi, plants, arthropods, and vertebrates across the dry-to-wet-season transition in a lowland tropical forest. All four assemblages shifted abruptly and nearly synchronously following rainfall late in the dry season, before the meteorological wet season began. These compositional transitions were largely independent of taxonomic richness and airborne DNA concentrations, indicating coordinated changes in detection patterns rather than simple seasonal gains in richness. Across all samples, we recovered more than 5,400 operational taxonomic units, including 2,148 fungi, 857 plants, 2,157 arthropods, and 239 vertebrates. We also recovered 949 full-length insect DNA barcodes, demonstrating compatibility with standard barcoding approaches and expanding opportunities for species discovery. Because the onset of the meteorological wet season showed no long-term trend at Barro Colorado Island, predicting future phenological and compositional shifts will require greater attention to dry-season rainfall variability and the cues that organisms track during this transition. Airborne environmental DNA offers a scalable, noninvasive framework for monitoring biodiversity and phenology across taxonomic groups.

## Introduction

Terrestrial biodiversity is declining across most major taxonomic groups, threatening ecosystem function and resilience [1, 2]. These losses are especially consequential in tropical forests, which harbor a disproportionate share of global biodiversity, much of it still undescribed [3-5]. Incomplete taxonomic knowledge obscures the magnitude of biodiversity change and limits our ability to understand how ecosystem processes respond to climate variability [4, 6, 7].

The seasonal dynamics of most tropical forest species are also poorly characterized. This gap limits predictions of how shifts in precipitation timing and magnitude will alter community composition and activity [8-10]. Seasonal climate can synchronize phenology, reproduction, and activity across taxa, coupling biological processes at ecosystem scales [11, 12]. In tropical ecosystems, precipitation may trigger coordinated responses among plants, animals, and microbes, with consequences for resource availability and species interactions [13-15]. Yet the environmental cues governing the timing and synchrony of these responses remain uncertain for most species [16].

Addressing these gaps requires biodiversity surveys that span taxonomic groups, reveal poorly documented diversity, and can be repeated frequently enough to resolve phenological change. Conventional survey methods capture only parts of this complexity and are often limited in breadth, comparability, or scalability [17-20]. Airborne environmental DNA (airborne eDNA), including DNA carried in spores, pollen, tissue fragments, and other particles, has the potential to characterize broad components of terrestrial biodiversity [21-23]. However, it remains unclear whether airborne eDNA can capture local signals from multiple taxonomic groups at the temporal resolution needed to quantify seasonal dynamics.

We examined seasonal patterns in airborne eDNA from Barro Colorado Island (BCI), Panama, a well-studied lowland tropical forest with a pronounced dry-to-wet-season transition that typically occurs in early May [24-26]. We sampled three sites weekly to test whether airborne eDNA assemblages of fungi, plants, arthropods, and vertebrates remained stable across the transition or changed before, during, or after the onset of rain. We also conducted spatially extensive campaigns in the late dry season and peak wet season to assess whether the weekly time series represented broader island-wide patterns. Finally, we generated standard barcode-length amplicons to evaluate the joint use of airborne eDNA for phenological monitoring and species discovery.

## Results and Discussion

### Characterizing seasonal transitions

Examining airborne eDNA assemblage profiles among weekly 24-hour samples from 3 sites for 23 weeks, we detected significant shifts in profile compositions that coincided with the onset of rains in mid-April. Fitting a double-breakpoint piecewise regression to principal coordinate analysis scores of Jaccard distances of airborne eDNA profiles, all four assemblage airborne eDNA profiles shifted synchronously near the onset of the wet season (Fig. 1). Assemblage profiles began shifting on day of year (DOY) 98 ± 4 (s.e.) for fungi, 98 ± 7 for plants, 100 ± 7 for arthropods, and 107 ± 7 for vertebrates. The dry-to-wet-season transitions of the fungal and plant assemblages lasted 23 d each, the arthropod assemblage profile transition lasted 37 d, and the shift in the vertebrate assemblage lasted 38 d.

**Figure 1.**
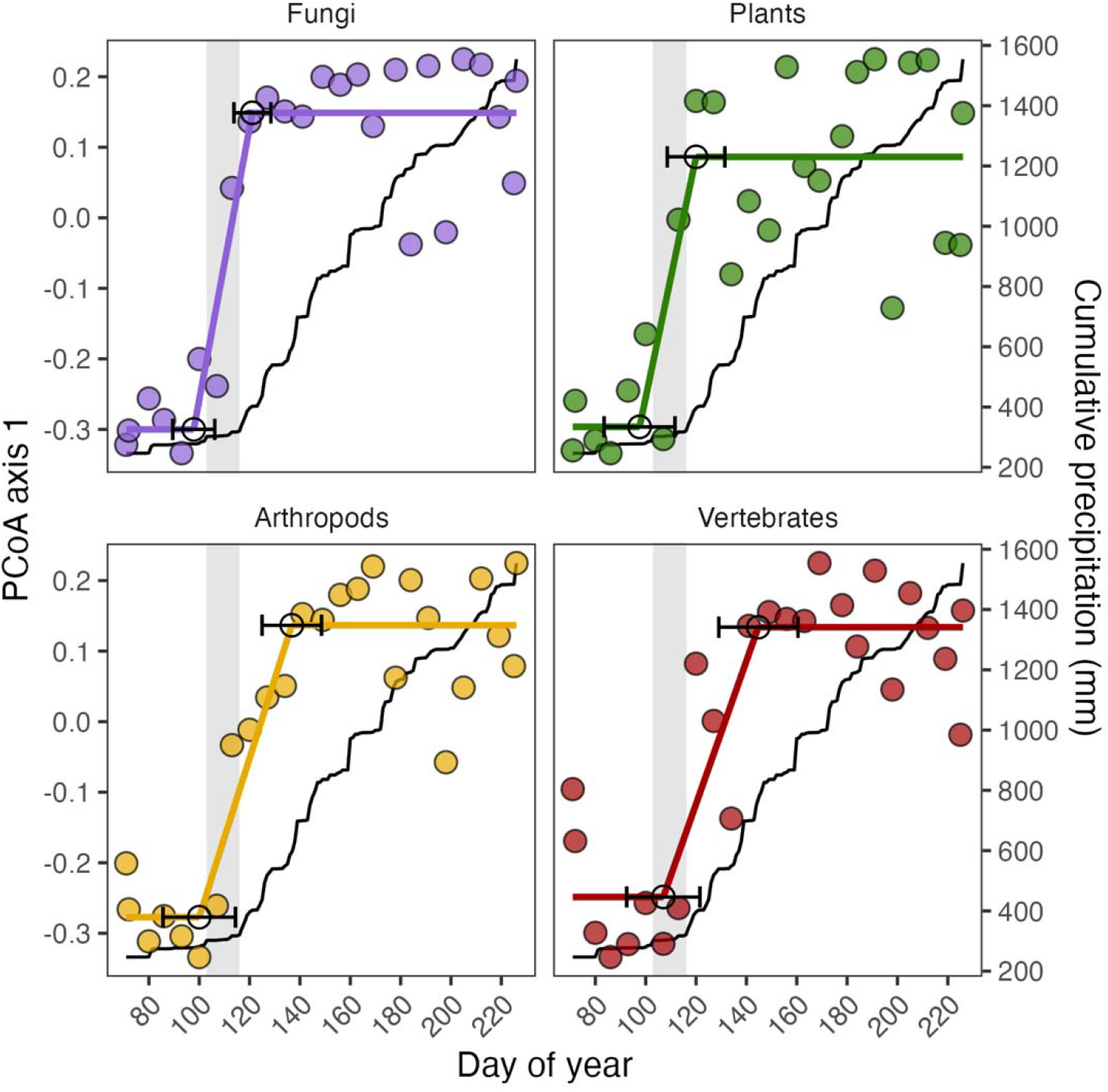
Seasonal shifts in airborne eDNA assemblage composition. First PCoA-axis scores for weekly samples and cumulative precipitation, with piecewise-regression breakpoints (open circles ± SE). Gray shading marks the interval between the initial precipitation event and meteorological wet-season onset.

The beginning of the shifts in airborne eDNA profiles coincide with a 24.5 mm precipitation event that fell from DOY 100-104 (Fig. S1), which was then followed by just 18 mm over the next 11 d. The onset of the meteorological wet season, defined by a breakpoint in the linear accumulation of precipitation over time (27) did not begin until approximately 2 weeks later (DOY = 116) when cumulative precipitation began to increase consistently at 10.5 mm d^-1^ vs. 1.9 mm d^-1^ from DOY 30-115 (Fig. S1, S2). The timing of the dry-to-wet season transition and dry-season precipitation amounts in 2025 were not atypical for this location. The 2025 breakpoint in cumulative precipitation was 7 d earlier than mean onset determined over the period from 1972-2024, but not atypically early (DOY = 123 ± 14 [s.d.]) (Fig. S2, S3). Also, significant precipitation events prior to the onset are also common: 21 of the years from 1972–2024 had precipitation events that exceeded 20 mm over a 3-d period at least 7 days prior to the onset of the wet season but after DOY 100.

The synchronous shifts in airborne eDNA profile composition were independent of temporal patterns in operational taxonomic unit (OTU) richness and DNA concentrations in air. Over the 23-week period from March to August, sample-level fungal OTU richness increased by 35% (*P* = 0.03) and plant OTU richness declined by 64% over this period (*P* < 0.001), but little of this shift occurred during airborne eDNA profile transition periods (Fig. 2). There was no significant linear change in richness for arthropod OTUs (*P* = 0.15) or vertebrate OTUs (*P* = 0.75) (Fig. 2). Similar to the richness patterns, we quantified marker gene DNA concentrations for each assemblage via quantitative PCR. Plant trnL DNA concentrations decreased over time, while fungal ITS, arthropod COI, and vertebrate 12S rRNA DNA concentrations were relatively unchanged over time (Fig. S4).

**Figure 2.**
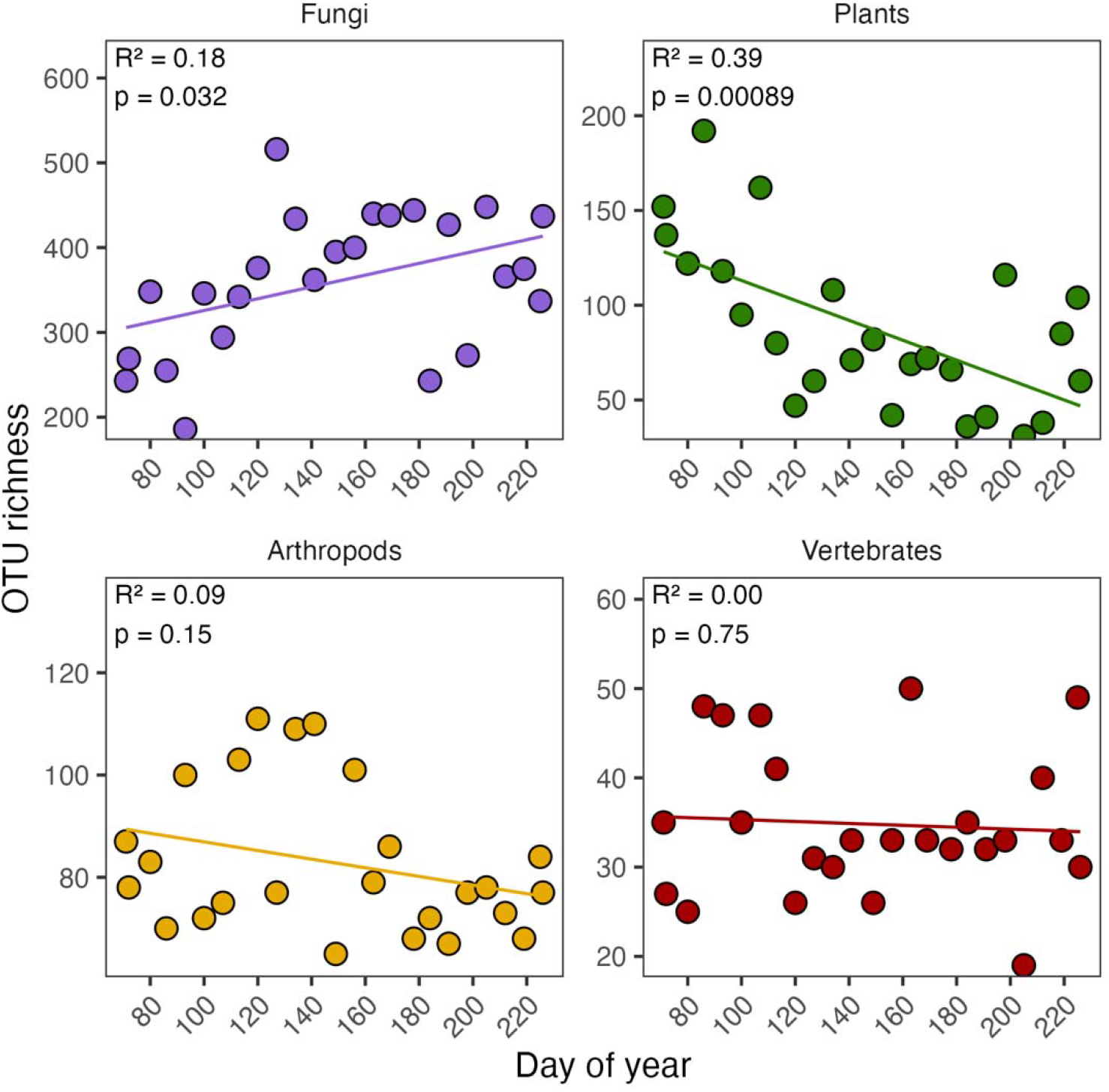
Temporal changes in airborne eDNA richness for each of four major groups sampled. OTU richness through time at the three sites sampled weekly, with linear regressions.

The shifts in assemblage profiles across the 23 weeks represented both decreases and increases in the prevalence of individual OTUs, whose identities provide ecological insights into the drivers of seasonal turnover of species’ DNA in the airborne eDNA profiles. Among fungal classes, Agaricomycetes (predominantly of the orders Agaricales and Hymenochaetales) increased from 61% to 81% of the total relative abundance of fungi from the dry to wet season, indicating that many litter- and wood-decaying fungi are strongly stimulated by sustained moisture. No other fungal classes changed more than 2% in mean relative abundance between seasons. Declines in plant richness were distributed across almost all plant orders, with the richness of OTUs of many predominantly dry-season flowering taxa, such as those in Sapindales and Lamiales, declining strongly after the onset of the wet season (Fig. S5). At the order level, the OTU richness of Porellales, Hypnales, Hookeriales, and Polypodiales (liverworts, mosses, and ferns) increased the most (Fig. S5). Among arthropods, Diptera (96% to 100%) and Cecidomyiidae (92% to 100%) were present in nearly all replicates both before and after the onset of the wet season, though a large number of OTUs from previously uncharacterized Cecidomyiidae taxa either decreased or increased in prevalence (Fig. 3). Trombidiformes (Arachnida) and Entomobryomorpha (Collembola) increased the most in prevalence between dry and wet seasons (10% to 39% and 8% to 33%, respectively), and Coleoptera (Insecta) and Ixodida (Arachnida) decreased the most (39% to 17% and 19% to 4%, respectively). Among vertebrates, many mammal OTUs declined while many bird OTUs increased in prevalence (Fig. 3). Among individual vertebrate species OTUs, declines in *Cuniculus paca* (lowland paca) and *Alouatta palliata* (howler monkey) prevalence were prominent and occurred with clear increases in the prevalence of ground-nesting *Baryphthengus martii* (rufous motmot) and *Amazona farinosa* (southern mealy amazon) (Fig. 3, Fig. S6). An overall reduction in the prevalence of mammals in airborne eDNA profiles over time may represent shifts in habitat use in response to rainfall-linked fruit and insect pulses (28, 29) though the observed patterns are unlikely to represent decreases in population size.

**Figure 3.**
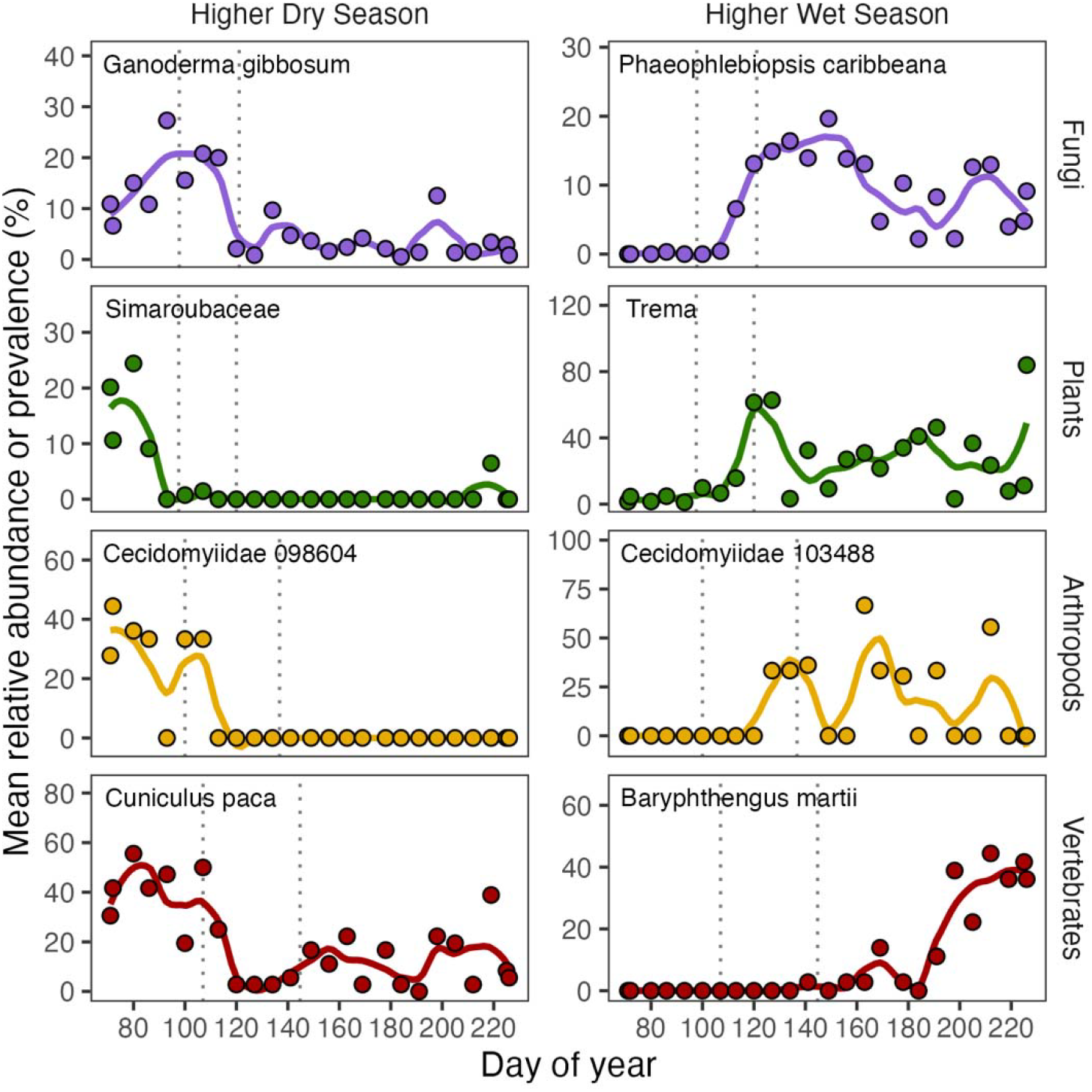
OTUs associated with the seasonal transition. Selected OTUs with higher relative abundance (fungi and plants) or prevalence (arthropods and vertebrates) before or after the compositional shift. Dotted lines mark estimated breakpoints in compositional shifts (see Fig. 1).

Although atmospheric conditions such as humidity and rainfall can influence the atmospheric fate and transport of airborne DNA (22), variation in transport, deposition, or degradation does not provide a parsimonious explanation for the coordinated, bidirectional, and taxon-specific shifts observed across four assemblages. Explaining these patterns through fate and transport alone would require coincident taxon-specific changes in particle association or relative persistence across independent genetic markers. Likewise, since the observed changes in total DNA concentrations or richness levels did not correspond to the timing of the shifts in assemblage compositions (Fig. 2), it is unlikely that the observed patterns are a direct result of rainfall associated removal of airborne DNA or DNA-containing particles. Instead, the observed patterns in assemblage composition are most consistent with seasonal differences in the production and aerosolization of biological material containing DNA.

### Seasonal campaigns

Weekly sampling was complemented with two, more spatially comprehensive airborne eDNA sampling campaigns, coinciding with the beginning and end of the weekly sampling effort discussed above. Each campaign consisted of filtering air for 24-hours at 27 sites each day for 2 days (30). Shifts in airborne eDNA profiles observed in weekly samplings at the three sites broadly parallel observed differences quantified with the more extensive seasonal campaigns. For example, campaign-scale sampling captured the high compositional turnover and the low overlap of taxa between seasons, even in assemblages where richness changed little through time (Fig. 4, Fig. 5). The observed richness values for the airborne eDNA profiles, both sample-level and total richness, were higher for fungi in the wet season, higher for plants in the dry season, and equivalent for arthropods and vertebrates between seasons (Fig. 4; Table S1,S2). Fungal DNA concentrations were 18% higher, on average, during the wet season than the dry season, plant DNA concentrations were 81% lower, and arthropod and vertebrate DNA concentrations were similar across the two seasons (Table S3, Fig. S7). Across the four assemblages, taxonomic turnover was high between the seasonal campaigns regardless of changes in profile richness, with only 25%, 35%, 13%, and 46% of OTUs shared between seasons for fungi, plants, arthropods, and vertebrates, respectively (Fig. 5). Differences in the abundance or prevalence of individual species between the two individual campaigns mirrored the weekly time course in many ways (Fig. 6). For example, among vertebrates, similar to the weekly samples, it was typically mammals such as *Cuniculus paca* (lowland paca), *Bradypus* sp. (three-toed sloths), and *Nasua narica* (white-nosed coati) that had higher detection rates during the dry season campaign than the wet season, while *Baryphthengus martii* (rufous motmot) and *Coragyps atratus* (black vulture) as well as *Amazona farinosa* (southern mealy amazon) had higher detection rates during the wet-season campaign (Fig. 6).

**Figure 4.**
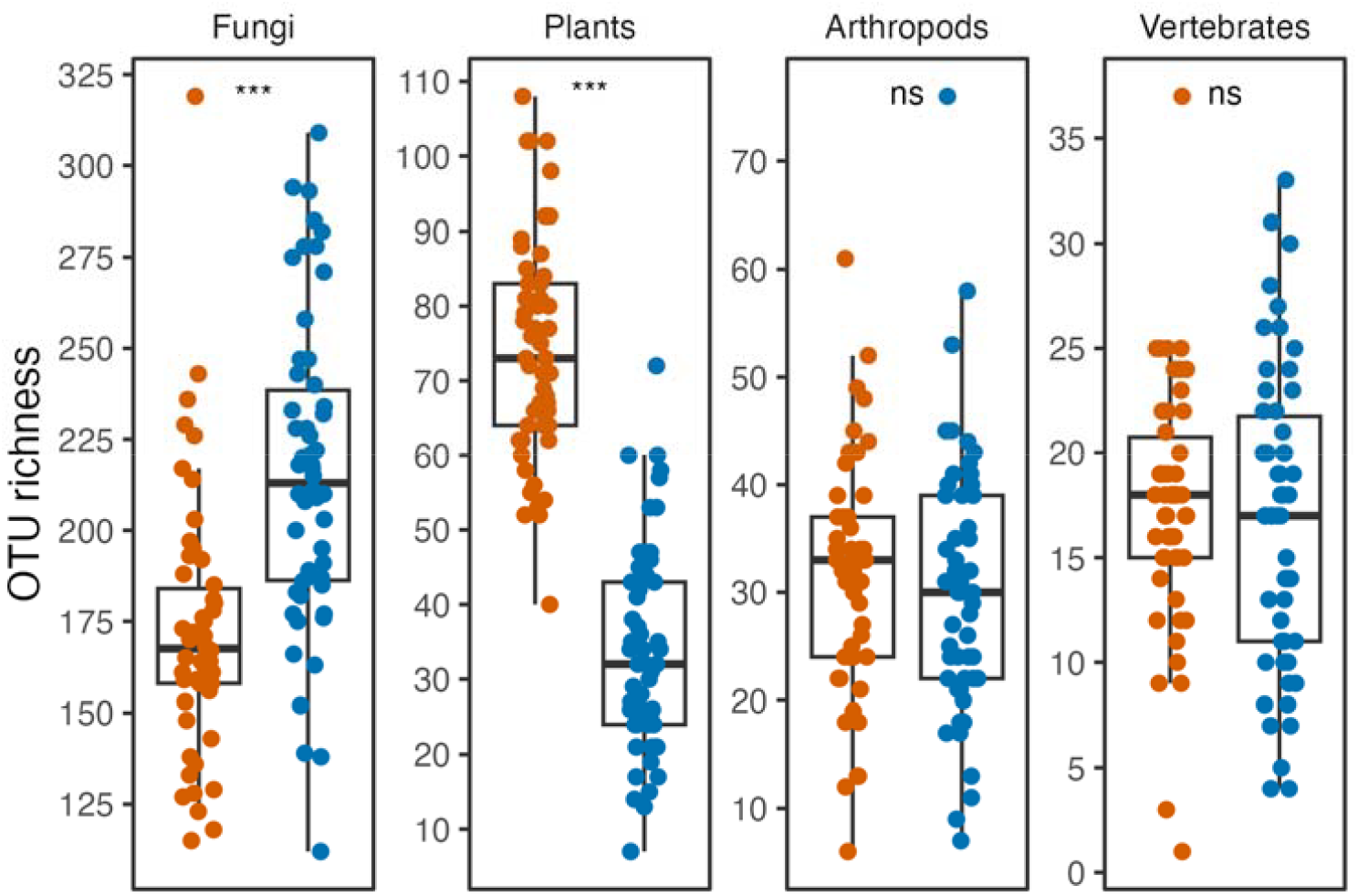
Seasonal differences in airborne eDNA OTU richness. Boxplots and site-level OTU richness during the dry-season (orange) and wet-season (blue) campaigns. ***P < 0.001; ns, not significant, P > 0.05.

**Figure 5.**
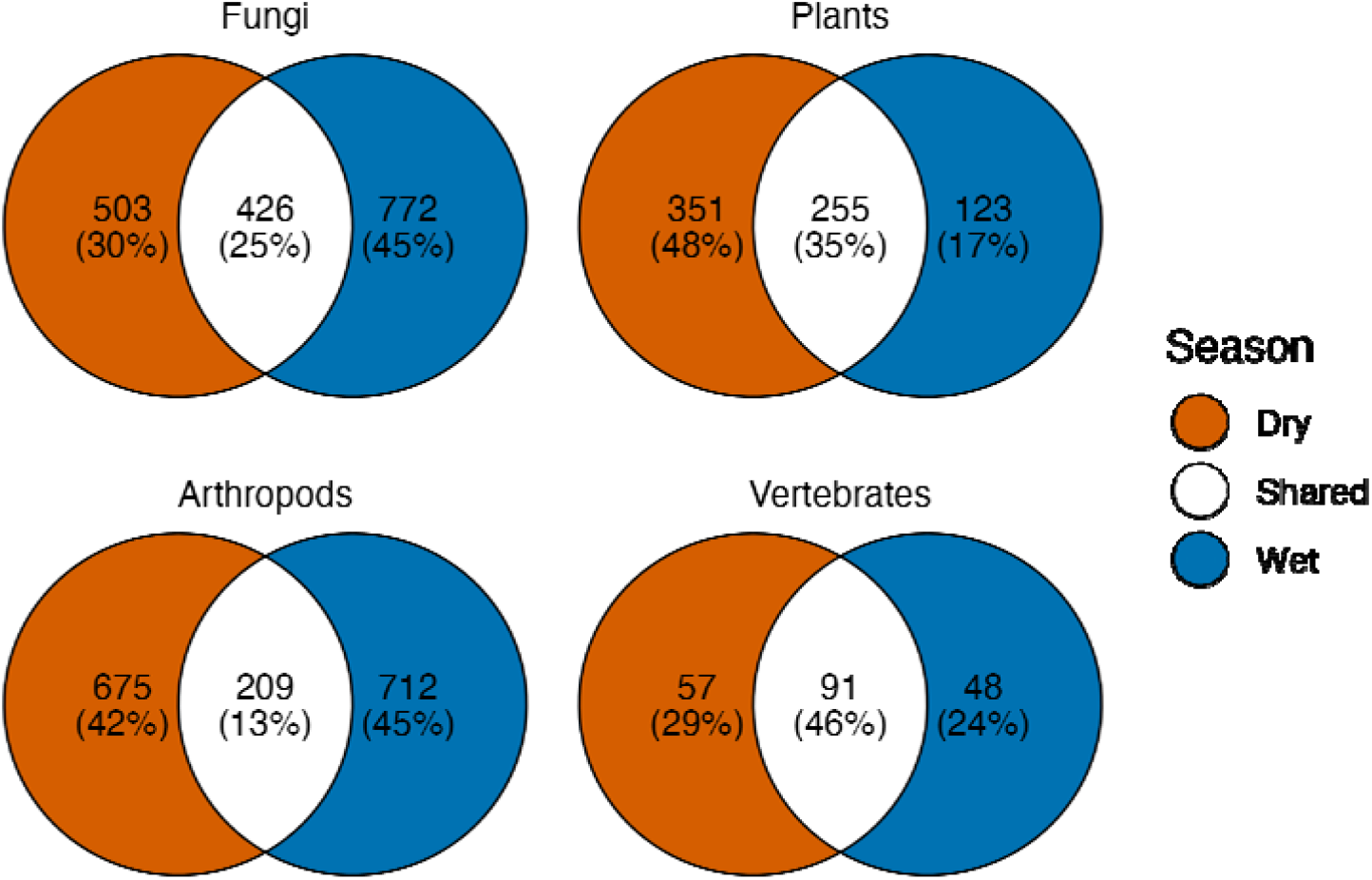
Seasonal overlap of airborne eDNA OTUs. Numbers and percentages of OTUs unique to the dry-season campaign (orange), shared between campaigns (white), or unique to the wet-season campaign (blue).

**Figure 6.**
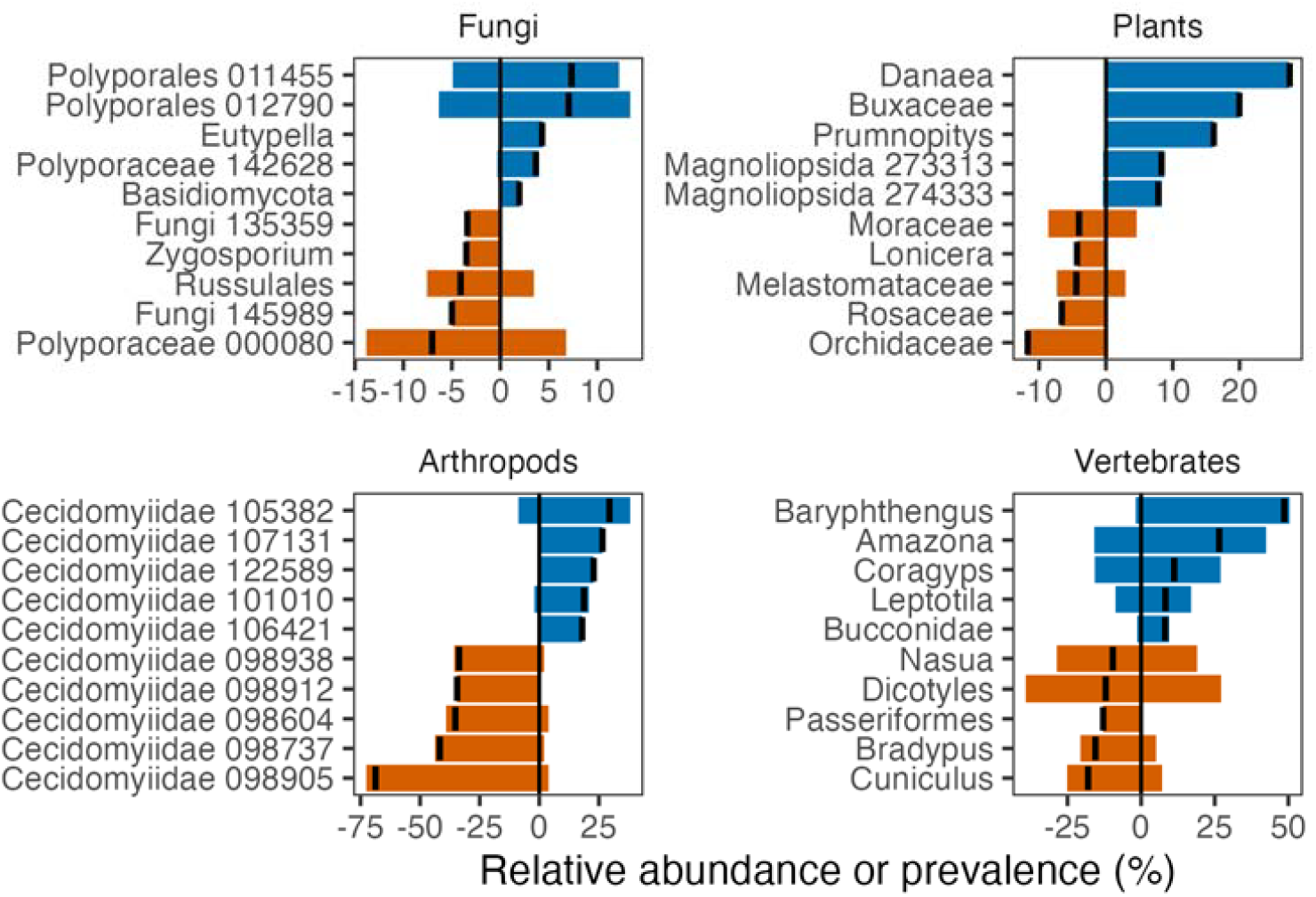
Taxa exhibiting the largest seasonal responses. The five largest increases and decreases in relative abundance (fungi and plants) or prevalence (arthropods and vertebrates) from the dry-to wet-season campaigns.

### Overall diversity patterns

When summed across the weekly samplings and the two campaigns, airborne eDNA assemblage profiles reflect the overall high diversity of the tropical forest, while also revealing diversity that has been poorly captured with traditional census techniques. Across the 171 samples acquired during the two campaigns and the weekly sampling, a total of 2148 fungal, 857 plant, 2157 arthropod, and 239 vertebrate OTUs were detected (Fig. 7). Among the 20 fungal classes detected, the greatest fungal richness was in Agaricomycetes (958), Sordariomycetes (478), and Dothideomycetes (192). For comparison, approximately 450 species of fungi have been documented from BCI using more traditional survey approaches (31). The 857 plant OTUs spanned 147 plant families, with Fabaceae (89 OTUs), Poaceae (54 OTUs), and Euphorbiaceae (41 OTUs) being the most diverse. By comparison, plant richness for the island is approximately 1300 species distributed across 166 families (32).

**Figure 7.**
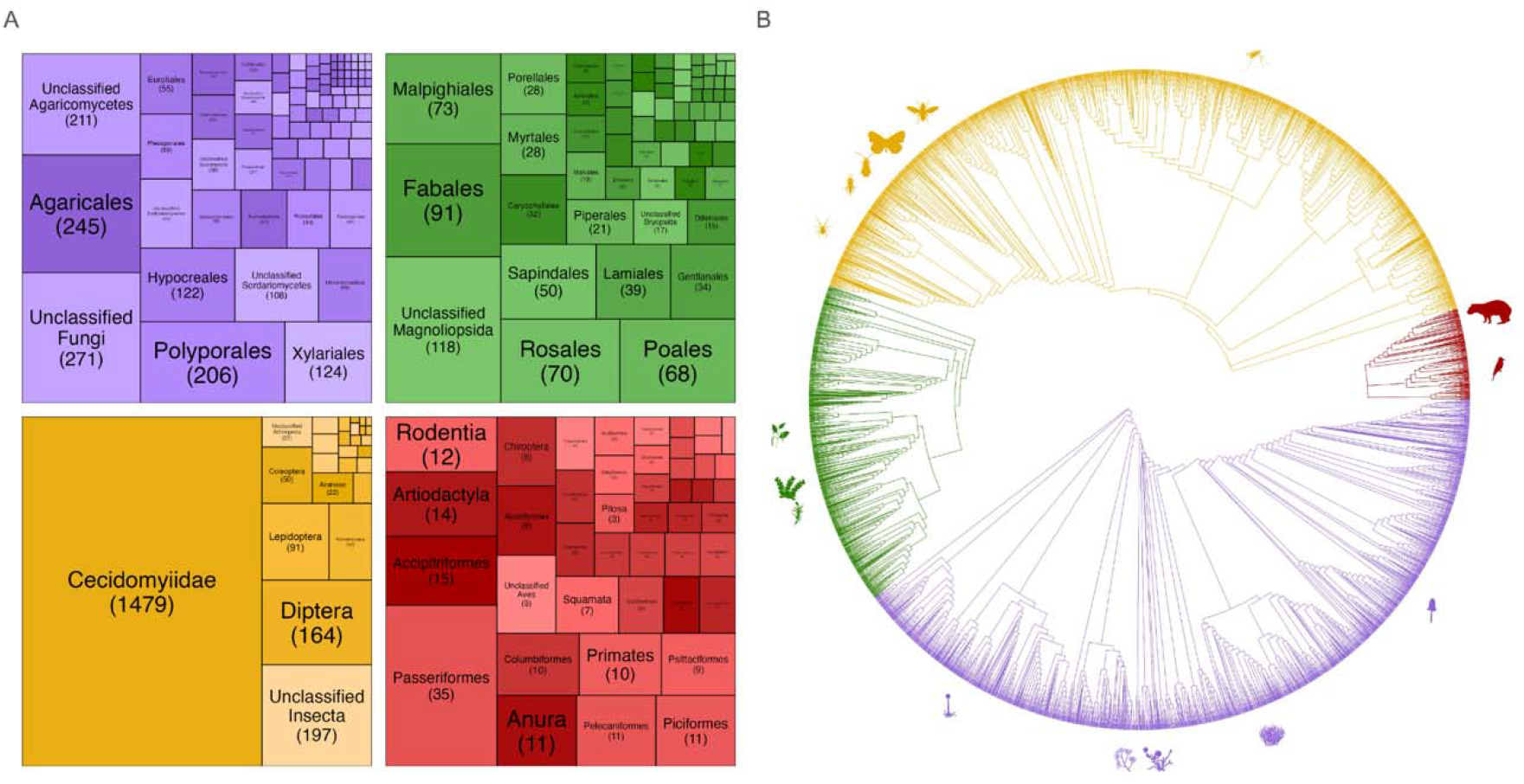
Taxonomic and phylogenetic diversity detected in airborne eDNA. (A) Treemaps of BCI OTU richness by order for fungi (purple), plants (green), arthropods (yellow), and vertebrates (red), combining weekly and seasonal-campaign samples. (B) Unrooted phylogenetic tree of OTUs across the four assemblages, using the same colors as panel A.

The 2157 arthropod OTUs spanned 6 classes and 22 orders. In total, 79% of all arthropod OTUs detected were Diptera (1709 OTUs), of which 87% were Cecidomyiidae (1479 OTUs), such that Cecidomyiidae alone accounted for 69% of all arthropod OTUs detected. Given the short adult lifespans characteristic of many Cecidomyiidae and repeated detections of many OTUs over time (Fig. S8), much of the detected Cecidomyiidae DNA likely originates from larval stages - potentially via frass or other indirect pathways - rather than adult activity alone. At the same time, this dominance of Cecidomyiidae does not imply an absence of other arthropod taxa in the airborne eDNA pool: each filter contained on the order of 10^8^ copies of arthropod mitochondrial DNA (Table S3), indicating that large quantities of DNA from non-Cecidomyiidae species may have been present but not detected due to amplification of more abundant species. Among the 31% of arthropod OTUs that were not Cecidomyiidae, families with the highest detected OTU richness were Tachinidae (2.36% of total arthropod OTUs, 51), Chironomidae (1.16%, 25), and Psychodidae (0.93%, 20), with 10.85% of arthropod OTUs (234) unclassified at the family level.

Of the 239 vertebrate OTUs, 144 OTUs were assigned to birds, 58 mammals, 17 fishes, 12 amphibians, and 8 reptiles. Known terrestrial vertebrate diversity for BCI includes at least 170 year-round resident bird species and another 120 non-resident bird species (33), 82 mammal species, approximately 130 reptile and amphibian species (34, 35). Of the 144 bird OTUs detected, 32 were exact matches to resident species while the rest were either likely residents that could not be confidently resolved to species with current reference databases or non-resident species that occasionally are present on BCI. Mammals included species from 24 families, including 13 rodents, 10 primates, 8 bats, 4 carnivores, and an armadillo. Amphibians were all anurans except for one caecilian (*Oscaecilia ochrocephala*), while reptiles spanned 6 families and included *Boa constrictor* and *Iguana iguana*.

OTU accumulation curves across samples indicate that weekly sampling accumulated taxa more rapidly than the short campaigns, reflecting the high turnover of many taxa in airborne DNA across the seasonal gradient. Integrating weekly sampling with the two campaigns yielded the highest cumulative richness (Fig. S9), demonstrating the value of combining temporally distributed sampling with intensive short surveys for biodiversity survey efforts. When all samples were aggregated, fungal richness was closest to saturation (20% additional taxa predicted), while plant and vertebrate richness showed intermediate gaps (40–45% additional taxa) (Fig. S9). In contrast, arthropods exhibited the largest difference between observed and estimated total richness (2157 vs. 5070), suggesting that >135% additional arthropod taxa could be recovered with further airborne eDNA sampling (Fig. S9).

### Detecting novel diversity

A closer examination of Cecidomyiidae DNA sequences highlights the potential for airborne eDNA to reveal substantial numbers of undocumented species as well as their phenologies. As of March 2025, the Barcode of Life Database (BOLD) contained 787 Cecidomyiidae sequence clusters (barcode index numbers, i.e. BINs) from Panama. In contrast, the BCI airborne eDNA surveys identified 1479 Cecidomyiidae COI OTUs, of which 80% (1183 OTUs) were novel, sharing <97% sequence similarity with Panamanian BOLD BINs. Cecidomyiidae is considered among the most diverse families of insects (36), and their dominance in the airborne eDNA assemblage suggests that their abundance and local richness are greatly underestimated by conventional sampling approaches (20). The novel OTUs were broadly and evenly distributed across the Cecidomyiidae phylogeny (Fig. 8A). Mean relative read abundance of airborne eDNA OTUs was only weakly correlated with sequence similarity to the closest BOLD BIN (*r* = 0.12, *P* < 0.001), indicating that novel OTUs were not restricted to taxa for which few reads were detected.

**Figure 8.**
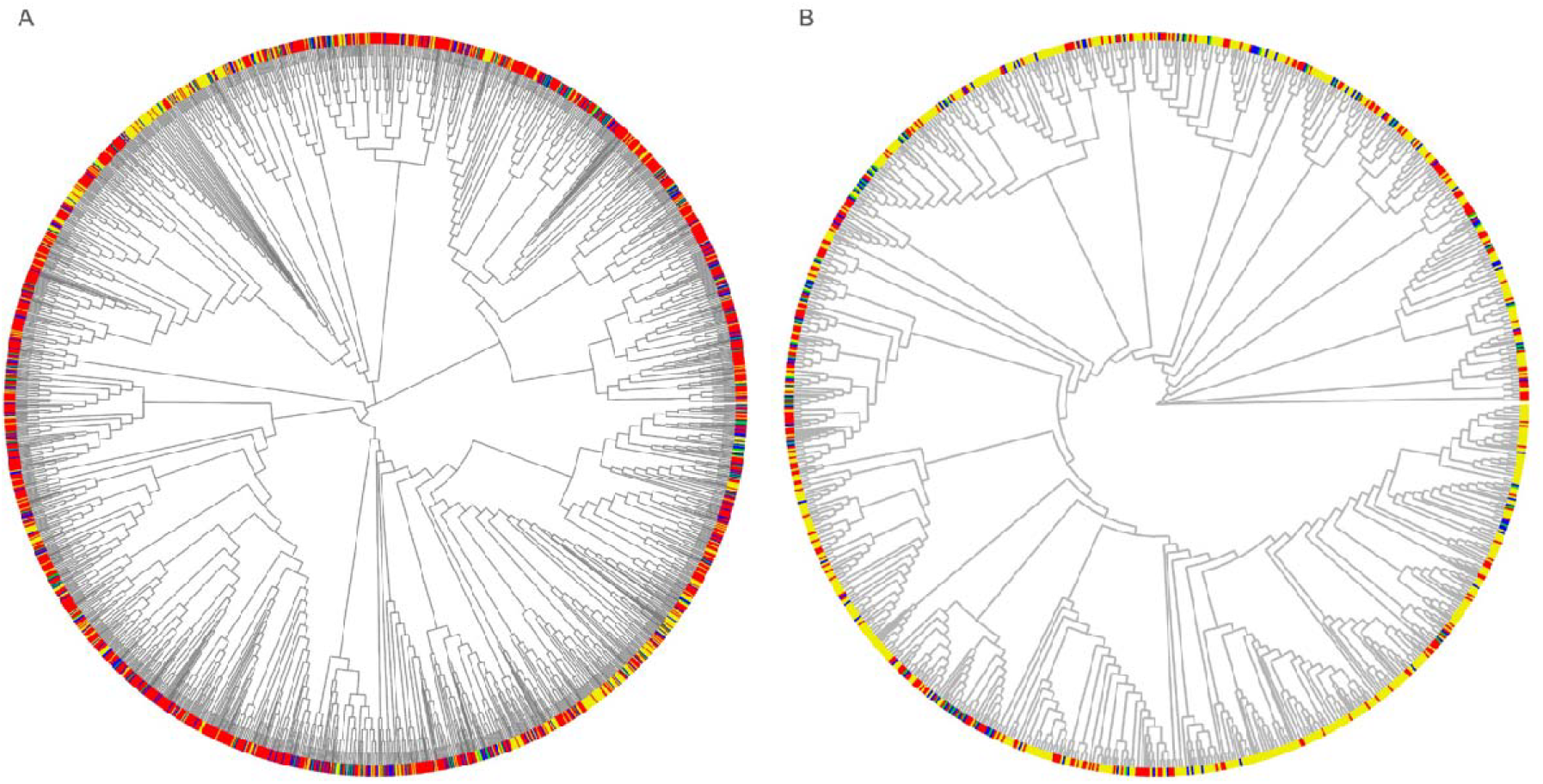
Phylogenetic distribution of Cecidomyiidae sequences from Barro Colorado Island. Trees compare (A) short-COI and (B) Folmer-COI OTUs with Panamanian BOLD barcode index numbers. BCI OTUs are colored by similarity to their closest BOLD record: below 97% (red) or at least 97% (blue). Panamanian reference sequences without an exact BCI match are colored below 97% (yellow) or from 97% to below 100% (green).

The COI fragments generated by our metabarcoding workflow are typically ∼157 bp, substantially shorter than the ∼658 bp Folmer COI barcode commonly used to describe metazoan species (37, 38). We conducted additional Nanopore sequencing with amplicons generated with the Folmer primer set on a single PCR replicate per sample to test whether standard barcode-length amplicons could be generated from airborne eDNA profiles to better characterize novel species. Across all BCI samples, 949 Folmer OTUs were generated and assignable to kingdom level. 837 (88%) of Folmer OTUs were from arthropods, 531 (56%) were from Diptera, and 409 (43%) were from Cecidomyiidae. Among Cecidomyiidae, 288 (70.4%) of Folmer OTUs were novel relative to Panamanian BOLD BINs, and these were similarly well distributed across the Cecidomyiidae phylogeny relative to BOLD accessions (Fig. 8B).

Patterns of novelty among fungi were comparable in magnitude to those observed for arthropods: 65% of the 2148 fungal OTUs detected at BCI did not match at >97% similarity to any sequences in the public UNITE database, potentially representing over a thousand previously unsequenced fungal species and providing further evidence that airborne eDNA analyses can be used for the detection of novel species.

### CONCLUSIONS

Overall, by sequencing airborne eDNA across a seasonal transition, we were able to document the reorganization of assemblage activities of fungi, plants, arthropods, and vertebrates that were synchronized by precipitation. These data provide strong evidence that the onset of the wet season synchronizes changes in activity for species across multiple assemblages in a tropical forest. Although there appear to be no significant trends in the onset of the meteorological wet season over the past 50 years at BCI (Fig. S3), we need to better understand how variation in the timing and pattern of precipitation onset affects seasonal transitions for different species. In general, sequencing airborne eDNA can quantify the presence and activity of a broad suite of species and monitor their changes in the phenology in the face of climate variability. As similar surveys extend across years, responses of species to interannual variation in climate should become clearer. Likewise, extending these surveys geographically will shed additional light on novel biogeographic patterns of assemblage composition and their phenological patterns.

## Methods

### Study site and air sampling

The study was conducted on Barro Colorado Island (BCI; 9.15 °N, 79.85 °W), a 1,542-ha lowland tropical forest reserve in Lake Gatun, Panama. The island lies 85-120 m above sea level, receives approximately 2,600 mm of rainfall annually, and has a mean daily maximum air temperature of 28.5 °C. It is almost entirely forested.

In March 2025, we established 27 airborne eDNA sampling sites along four trails: four on Snyder-Molina, nine on Shannon, eight on Barbour Lathrop, and six on Fairchild. Sites were located in forest, typically secondary forest less than 100 years old, spaced approximately 100 m apart, and positioned within a few meters of a trail.

For air sampling, 0.7-A, 120-mm fans were mounted in custom brackets and strapped to trees approximately 1.5 m above the ground. Each fan was powered by a 17-Ah, 12-V lead-acid battery. A custom filter card was secured in front of the fan with a plastic grate. Each card contained two 10-cm^2^ windows holding MERV 13-rated polyester filter material. Laboratory tests indicated an airflow of approximately 36 m^3^ h^−1^ with filters installed.

Sampling comprised two island-wide campaigns and weekly collections between them. The dry-season campaign began on 11 March 2025 and the wet-season campaign began on 12 August 2025. During each campaign, filters were deployed on two consecutive days. Mean filtration duration was 23.1 hours in the dry season (range, 21.2-25.1 hours) and 23.7 hours in the wet season (range, 22.4-24.5 hours). Between campaigns, filters were collected weekly after 24 hours at three sites (BL2, DF2, and SM4). Over 21 weeks, this generated an additional 63 samples. On 10 April, a 10-cm^2^ plastic cover extending approximately 5 cm beyond the fan was added above each filter to reduce direct rain exposure; all subsequent samples used this cover.

At collection, each filter was placed in a polyethylene bag with 5 g of silica desiccant and stored at 4 °C. Samples were transported to the Jonah Ventures laboratory in Boulder, Colorado, USA, on 15 March, 15 May, and 15 August 2025 and then stored frozen until extraction.

### Sample processing and sequencing

DNA was extracted from one of the two windows on each filter card using the Omega Bio-tek Mag-Bind Universal Pathogen Core Kit (M4030-01) according to the manufacturer’s protocol. Extractions were performed on a Hamilton STARlet running Venus 4. Approximately 300 µL of lysate was processed, and DNA was eluted in 100 µL of elution buffer and stored at -20 °C. We also extracted DNA from six filter cards that were not used for field sampling with these extraction blanks included in all downstream processing of the samples to check for introduction of potential contaminants.

Five marker regions were amplified from each genomic DNA sample: nuclear ITS for fungi (mean, 262 bp), chloroplast trnL for plants (145 bp), mitochondrial short COI (157 bp) and Folmer COI (656 bp) for arthropods, and mitochondrial 12S rRNA for vertebrates (395 bp). Primer sequences and thermal cycling conditions are provided in Table S4. Forward and reverse primers included 5^′^ adaptor sequences for subsequent indexing and sequencing.

One DNA aliquot per sample was amplified for ITS, trnL, and Folmer COI; 12 replicate aliquots were amplified for short COI and 12S rRNA. Each 25-µL reaction contained 12.5 µL Promega PCR Master Mix (M5133), 0.5 µL of each 10-µM primer, 1.0 µL genomic DNA, and 10.5 µL nuclease-free water. Amplification conditions are listed in Table S4.

Amplicons were treated with Exonuclease I and shrimp alkaline phosphatase (New England Biolabs M0293L and M0371L) for 30 minutes at 37 °C, followed by enzyme inactivation for 5 minutes at 95 °C. A second PCR added dual 10-bp multiplexing indices. Reactions contained Promega PCR Master Mix, 0.5 µM of each primer, and 2 µL of cleaned first-round product. Cycling consisted of 3 minutes at 95 °C followed by eight cycles of 30 seconds at 95 °C, 30 seconds at 55 °C, and 30 seconds at 72 °C. Indexed amplicons were purified and normalized with Cytiva SpeedBead magnetic carboxylate-modified particles (45152105050250) using a bead-based protocol [39]. Five microliters of each normalized sample were pooled. Libraries were prepared with the Oxford Nanopore Technologies Ligation Sequencing Kit V14 (SQK-LSK114) and sequenced on seven MinION flow cells using a GridION, generating 145.61 million reads.

Raw Nanopore data were basecalled from POD5 to FASTQ format with Dorado v0.7.1 and the super-high-accuracy <u>dna_r10.4.1_e8.2_400bps_sup@v5.0.0</u>model. Cutadapt v3.4 [40] was used to remove outer adapters and orient reads in the 5’ to 3’ direction. Reads were demultiplexed with pheniqs v2.1.0 [41], allowing no more than one mismatch in each paired 10-bp index. Cutadapt was then used to remove gene-specific primers and retain amplicons within marker-specific length ranges.

Exact sequence variants (ESVs) were identified by the UNOISE3 algorithm implemented in VSEARCH [42]. Candidate ESVs were required to occur at least four times across all samples and to have a maximum expected error below one base. Putative chimeras were removed with uchime3. Final read counts were obtained by mapping unfiltered reads to ESVs with usearch_global at a minimum identity of 95%. Taxonomy was assigned with a custom best-hit algorithm using UNITE for ITS, NCBI GenBank for trnL and 12S rRNA, and BOLD for COI. Exhaustive semiglobal pairwise alignments were performed with VSEARCH. Consensus taxonomy was based on all 100% matches or all references within one percentage point of the top match, and a taxonomic rank was accepted when more than 90% of top hits agreed.

We retained ESVs with at least 85% similarity to target phyla and conducted a second de novo chimera screen with uchime_denovo in VSEARCH using an abskew of 2. Marker-specific curation then removed trnL matches to temperate taxa unlikely to occur near BCI, including *Quercus* and *Pinus*; non-arthropods and Branchiopoda from COI; and human and domestic-animal sequences (*Canis, Felis, Equus, Sus*, and *Gallus*) from 12S rRNA. Sequencing depth was standardized within each assemblage by rarefying each sample to the minimum retained depth or 1,000 reads, whichever was greater. Rarefaction was performed with rrarefy in vegan and separately for BCI-only and regional comparisons. Rarefied ESVs were clustered at 97% identity with cluster_fast in VSEARCH to produce OTUs approximating species-level groups [37, 42]. The centroid taxonomy was assigned to each OTU. For 12S rRNA, assignments were compared with known species lists and multiple OTUs assigned to the same species were collapsed. Extraction controls contained only human and domestic-dog 12S rRNA sequences and were excluded from further analysis.

### Data analysis

We analyzed daily precipitation records from BCI for 1972-2025 using prorated manual measurements from the clearing near BCI headquarters [43]. Daily cumulative precipitation was calculated beginning on DOY 30. Analyses ended on DOY 270, except in 1986 and 1987, when they ended on DOY 225 because of anomalous late-season rainfall associated with a strong El Niño event. The year 2020 was excluded because precipitation records were incomplete. Wet-season onset was estimated by fitting a discontinuous piecewise regression to cumulative precipitation [27]. Candidate breakpoints between DOY 30 and 165 were evaluated with a one-breakpoint model that allowed accumulation rate to differ between dry- and wet-season segments. The breakpoint maximizing model fit, subject to minimum data requirements on each side, was retained.

For weekly time-series analyses, read counts were converted to presence or absence. Jaccard dissimilarities were ordinated with principal coordinate analysis (PCoA), and scores on the first axis were regressed against DOY 71-226 with a two-breakpoint piecewise linear model. When initial or terminal slopes did not differ significantly from zero, they were constrained to zero for visualization. Candidate breakpoint pairs were evaluated on a five-day grid spanning the first and second halves of the time series, respectively. Models were compared with the small-sample corrected Akaike information criterion, and the model with the lowest AICc was selected.

For each assemblage, we summarized OTU relative abundance (ITS and trnL) or prevalence across PCR replicates and samples (COI and 12S rRNA) before the first breakpoint and after the second breakpoint. We displayed the OTU with the largest decrease and the OTU with the largest increase for each assemblage.

For seasonal campaign comparisons, OTU counts were summed across the two sampling days at each site. OTU richness was compared between seasons with independent two-sample t-tests. Seasonal overlap was visualized with ggVennDiagram, and the five taxa with the largest positive and negative seasonal differences in relative abundance or prevalence were displayed with mirror plots in ggplot2.

Treemaps summarized OTU richness by order or family. For each marker, unique OTU sequences were aligned with MAFFT v7.526 using the auto option. Maximum-likelihood phylogenies were inferred in R with phangorn from a BIONJ starting tree based on maximum-likelihood distances and optimized under a GTR model with gamma-distributed rate heterogeneity and invariant sites. Assemblage trees were grafted onto a scaffold tree with ape and combined for visualization with ggtree; branch lengths and basal branches were omitted from the display.

Sampling completeness was evaluated for each assemblage in four datasets: all sampling events combined, weekly sampling, the dry-season campaign, and the wet-season campaign. Asymptotic richness was estimated with the Chao1 estimator in iNEXT, and incidence-based rarefaction and extrapolation curves were generated to 95% of the estimated asymptote.

Cecidomyiidae COI sequences were compared with Panamanian BOLD records annotated as COI-5P and downloaded in March 2025. Reference records were dereplicated and trimmed to the short-COI or Folmer target region with Cutadapt and similarity-guided extraction in VSEARCH. Reference and study sequences were aligned with MAFFT, and pairwise similarities were calculated from maximum-likelihood distances. Each OTU was assigned to the closest BOLD sequence and categorized as 100%, at least 97% but below 100%, or below 97% similarity. One reference sequence per barcode index number was retained for maximum-likelihood phylogenetic analysis under a GTR model.

Quantitative PCR was used to estimate assemblage-specific marker-gene concentrations across time and between seasons. Raw copy numbers were multiplied by the proportion of sequencing reads retained for the corresponding marker and sample. Seasonal differences were tested separately for each assemblage with two-sample t-tests on log-transformed concentrations. Weekly dynamics were summarized with site- and assemblage-level time series and simple linear regressions.

## Acknowledgements

We thank Lovisa Duck, Adalberto González, Manuel García, and Donato Jimenez for assistance with sample collection, and Lizzie Wolkovich and John Battles for comments on the manuscript.

## Author contributions

J.M.C. and N.F. conceived the study. J.M.C., N.F., K.S., N.S., M.R., D.L., and J.D. developed the methodology. J.M.C., N.F., K.S., J.D., and D.L. conducted the investigation. J.M.C., N.S., and D.L. performed the formal analysis. J.M.C., N.F., N.S., and D.L. curated the data. J.M.C. and N.F. administered the project. J.M.C. wrote the original draft. J.M.C., N.F., K.S., and N.S. reviewed and edited the manuscript. J.M.C. and N.S. prepared the visualizations.

## Data availability

Processed data supporting the results are provided in the Supplementary Data workbook. Repository accession numbers for raw sequence data and analysis code will be added before publication.

