## Supplemental Figures and Tables for "Airborne DNA reveals synchronized responses of tropical forest assemblages to precipitation"

**List of Supplementary Materials**
Figs. S1 to S9
Tables S1 to S4
Supplementary Data

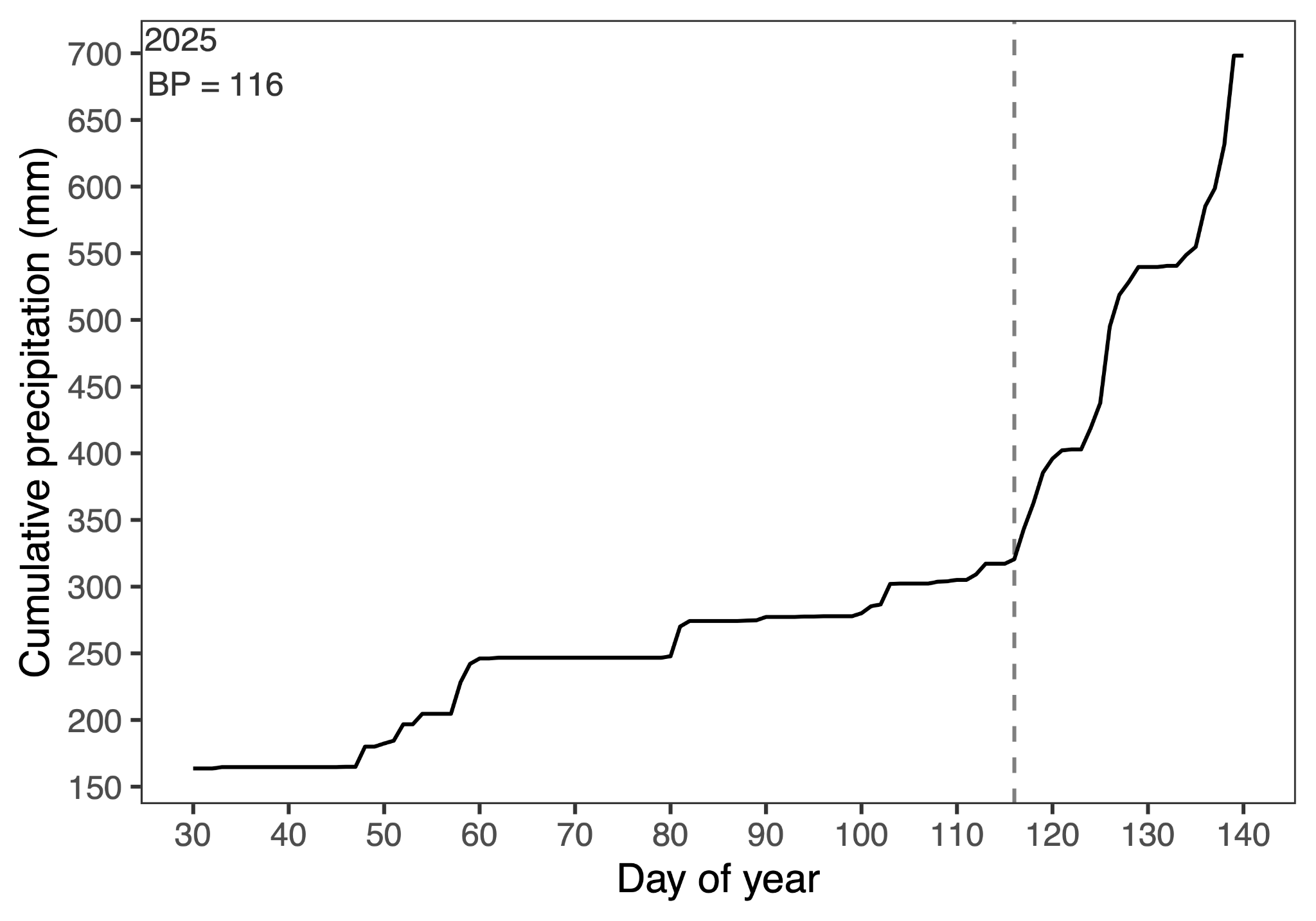

**Fig. S1.** Cumulative precipitation for 2025 highlighting the time around the transition from the dry to wet season.

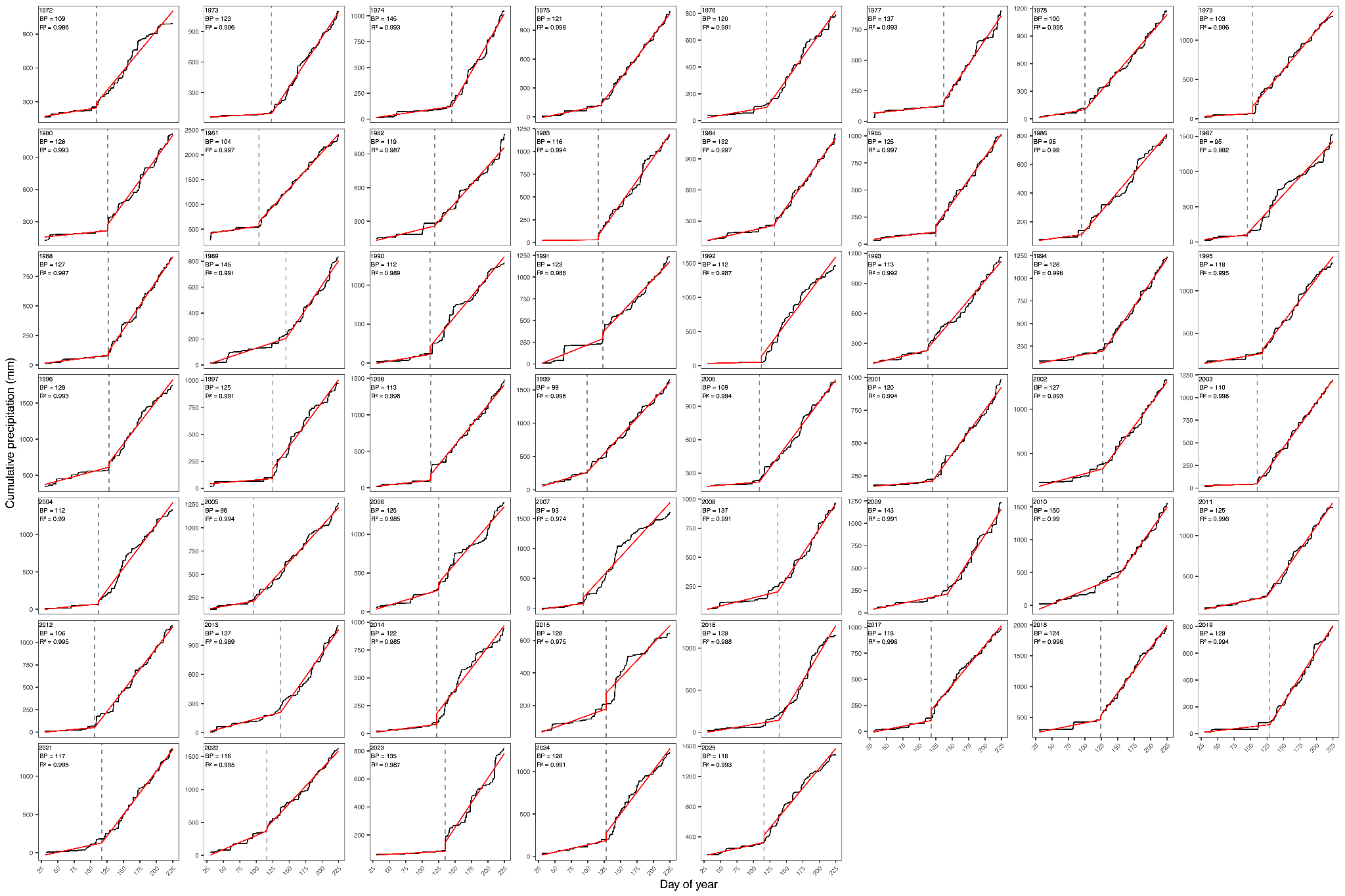

**Fig. S2.** Cumulative precipitation beginning day of year 30 for each year from 1972 – 2025. Shown are piecewise regression results (red lines). Breakpoints equivalent to the onset of the wet season are shown with vertical dashed lines.

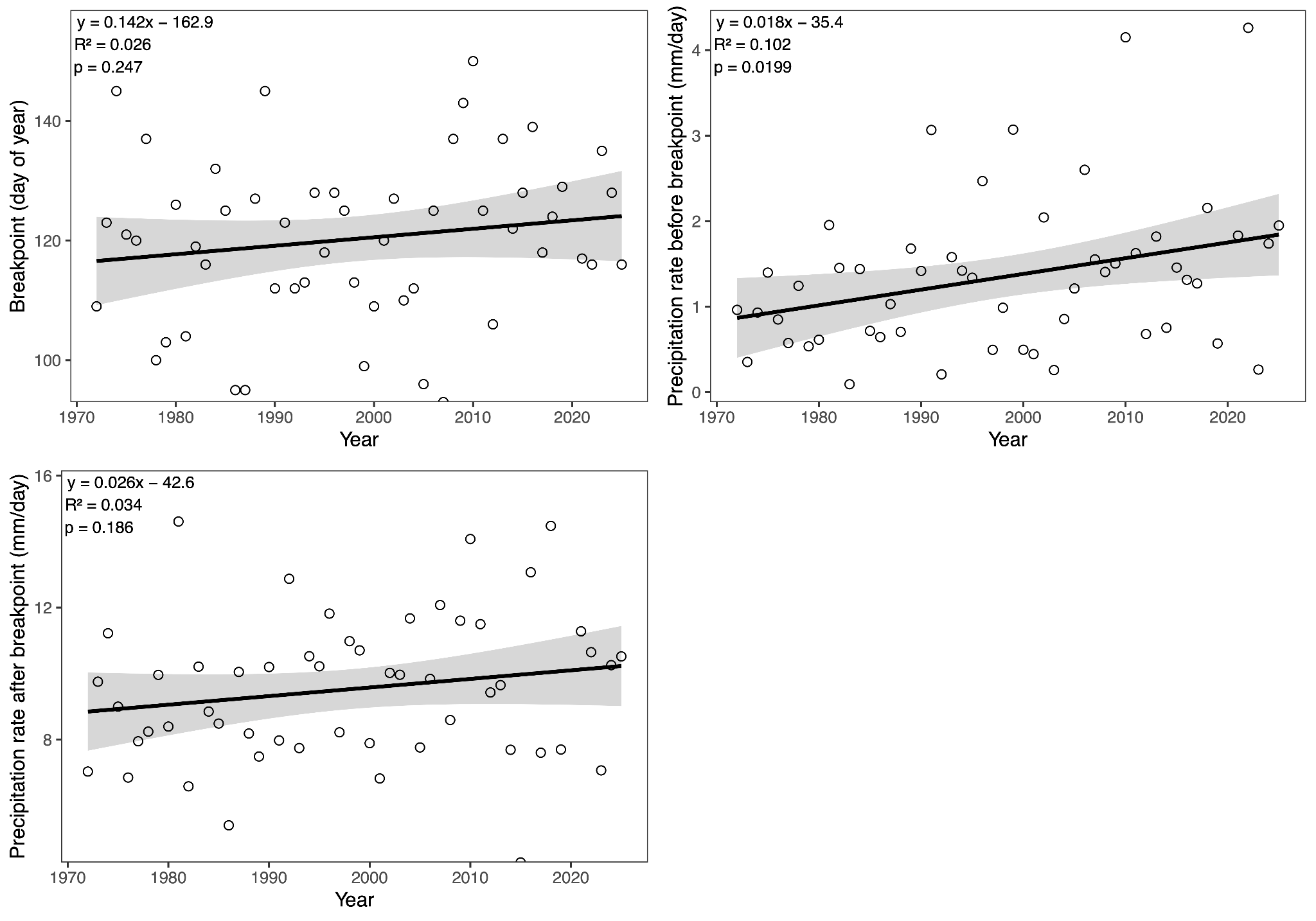

**Fig. S3.** Bivariate plots of year (1972 – 2025) vs. coefficients from yearly piecewise regressions of cumulative precipitation for (A) breakpoints of the onset of the wet season, (B) rate of cumulative precipitation in the dry season, and (C) rate of cumulative precipitation after the onset of the wet season. Linear regression results are included for each panel.

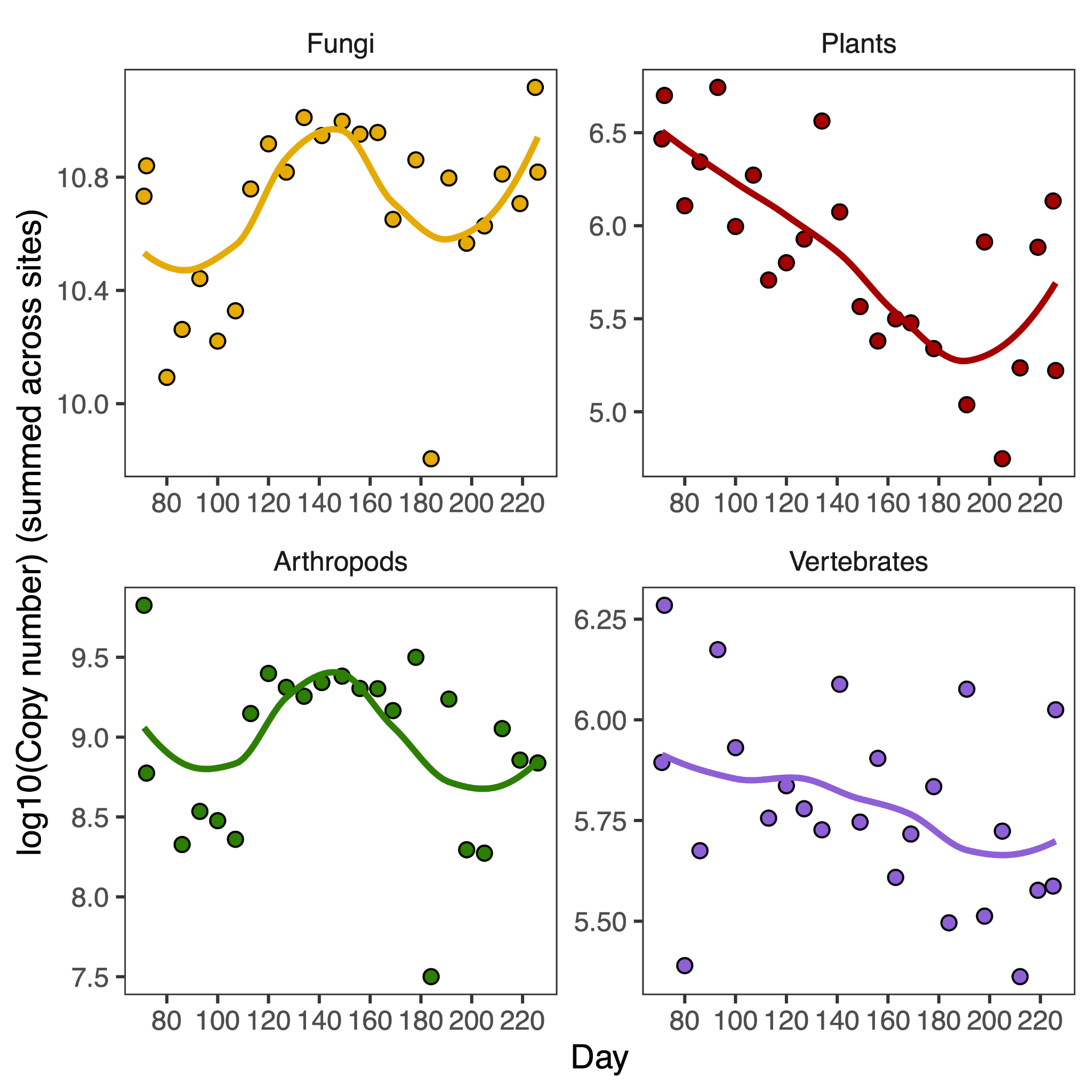

**Fig. S4.** Changes in target copy numbers across BCI weekly samples for each of the four assemblages as determined with qPCR. Linear regressions are shown.

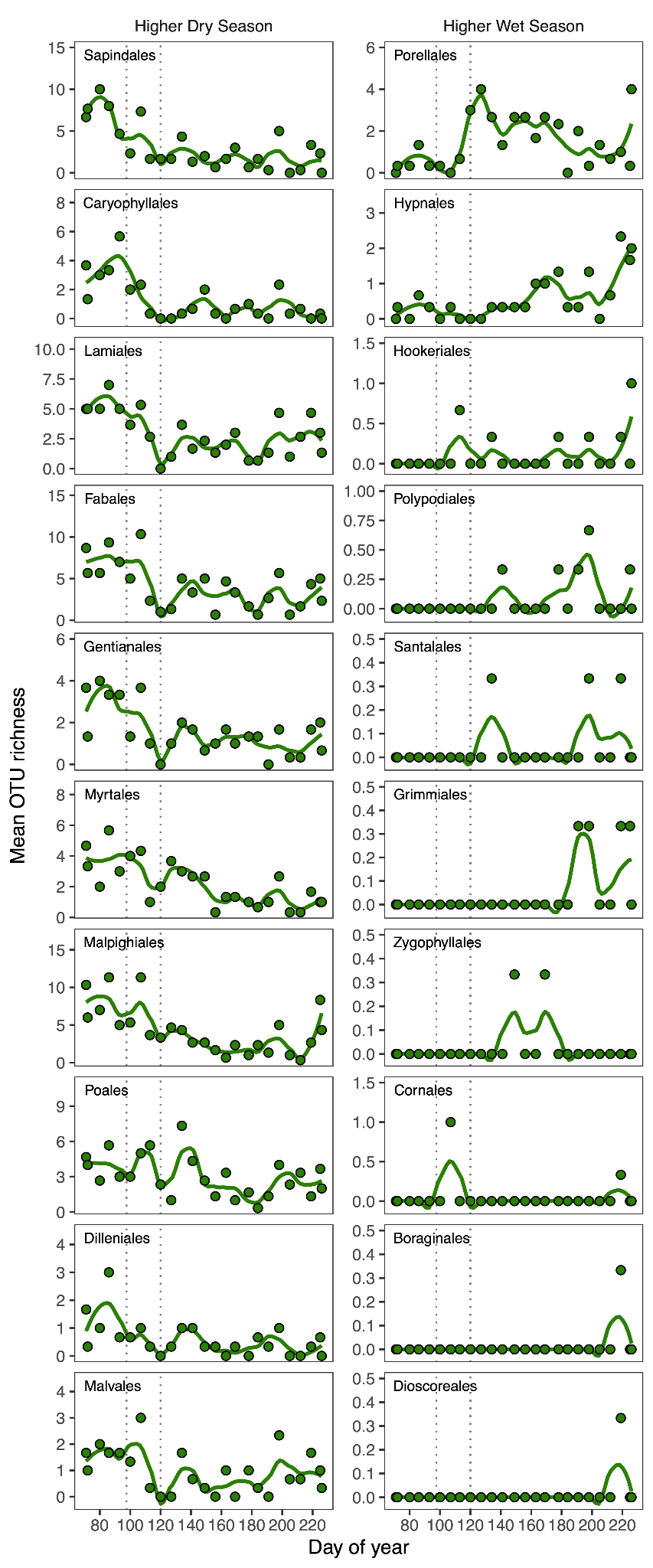

**Fig. S5.** Richness of plant orders across BCI weekly samples for the top 10 orders with greater richness in the dry season (left) and wet season (right).

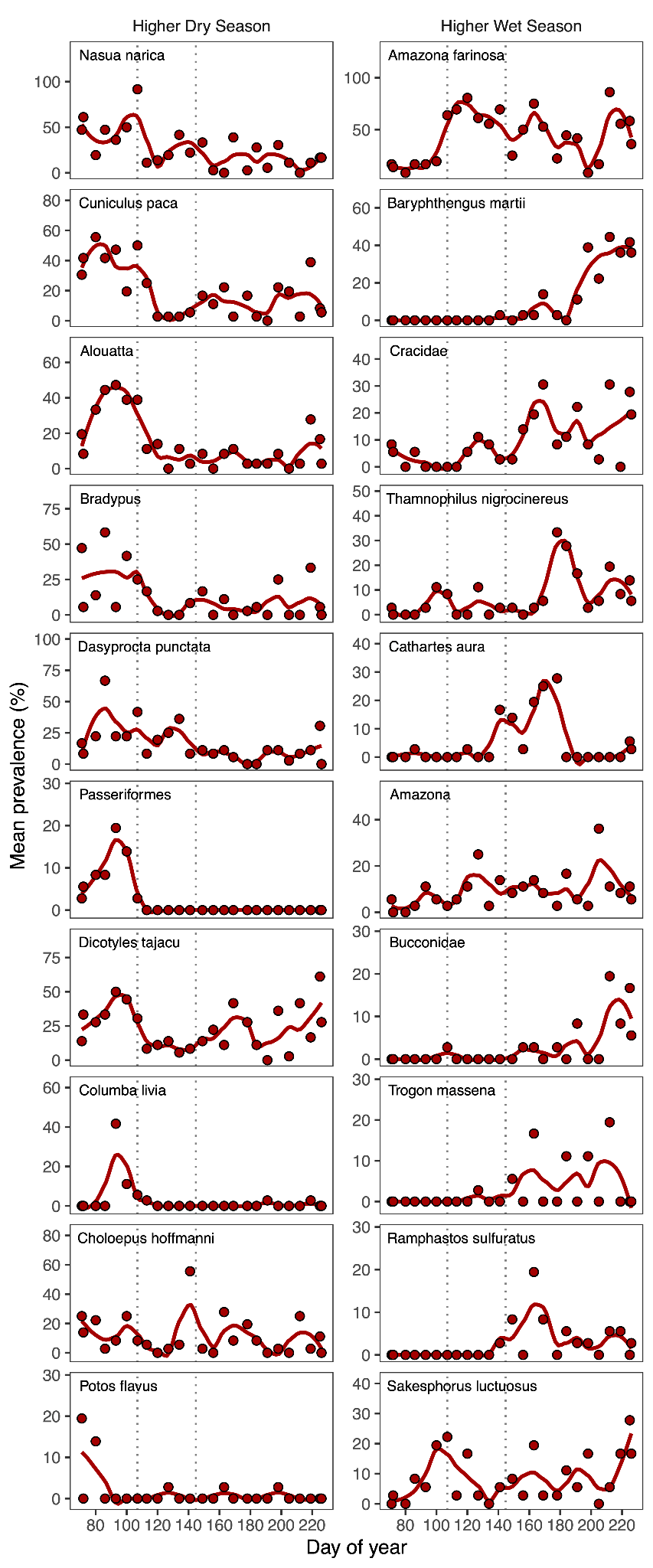

**Fig. S6.** Prevalence of vertebrate species across BCI weekly samples for the top 10 vertebrate OTUs with greater prevalence in the dry season (left) and wet season (right).

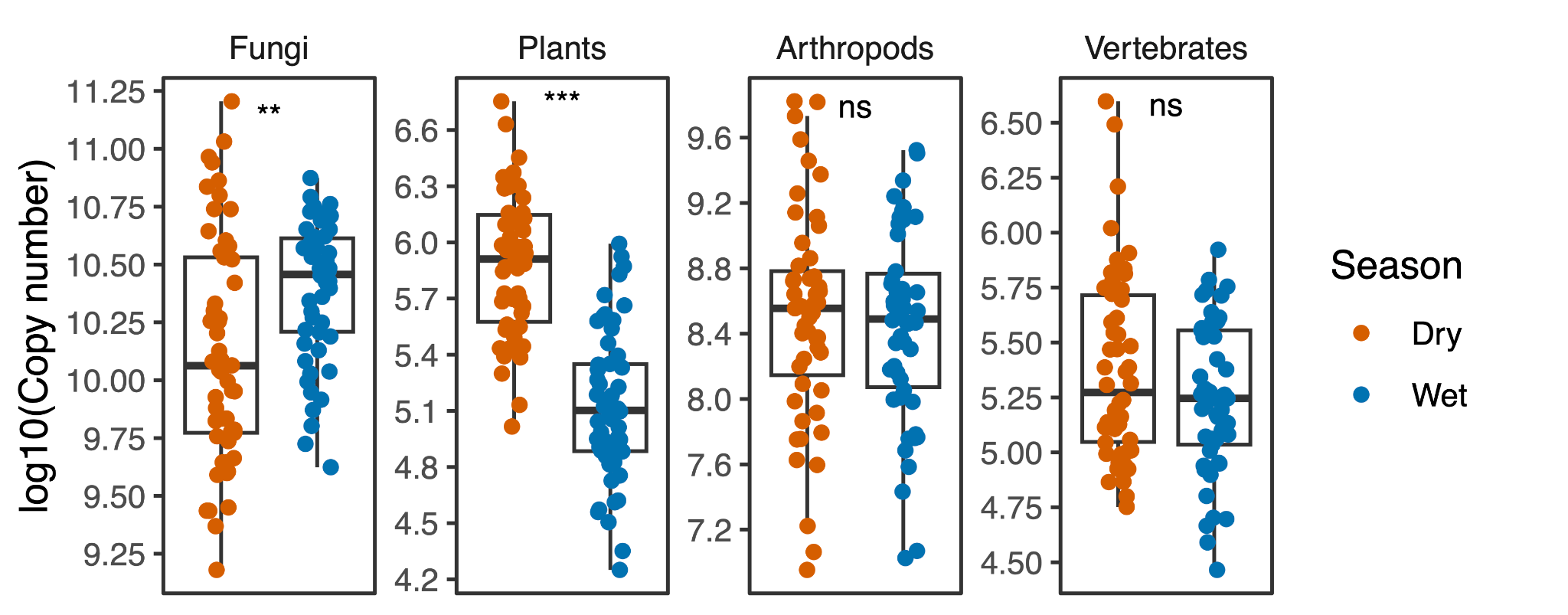

**Fig. S7.** qPCR copy number comparisons for all four assemblages between dry and wet seasonal sampling campaigns in BCI.

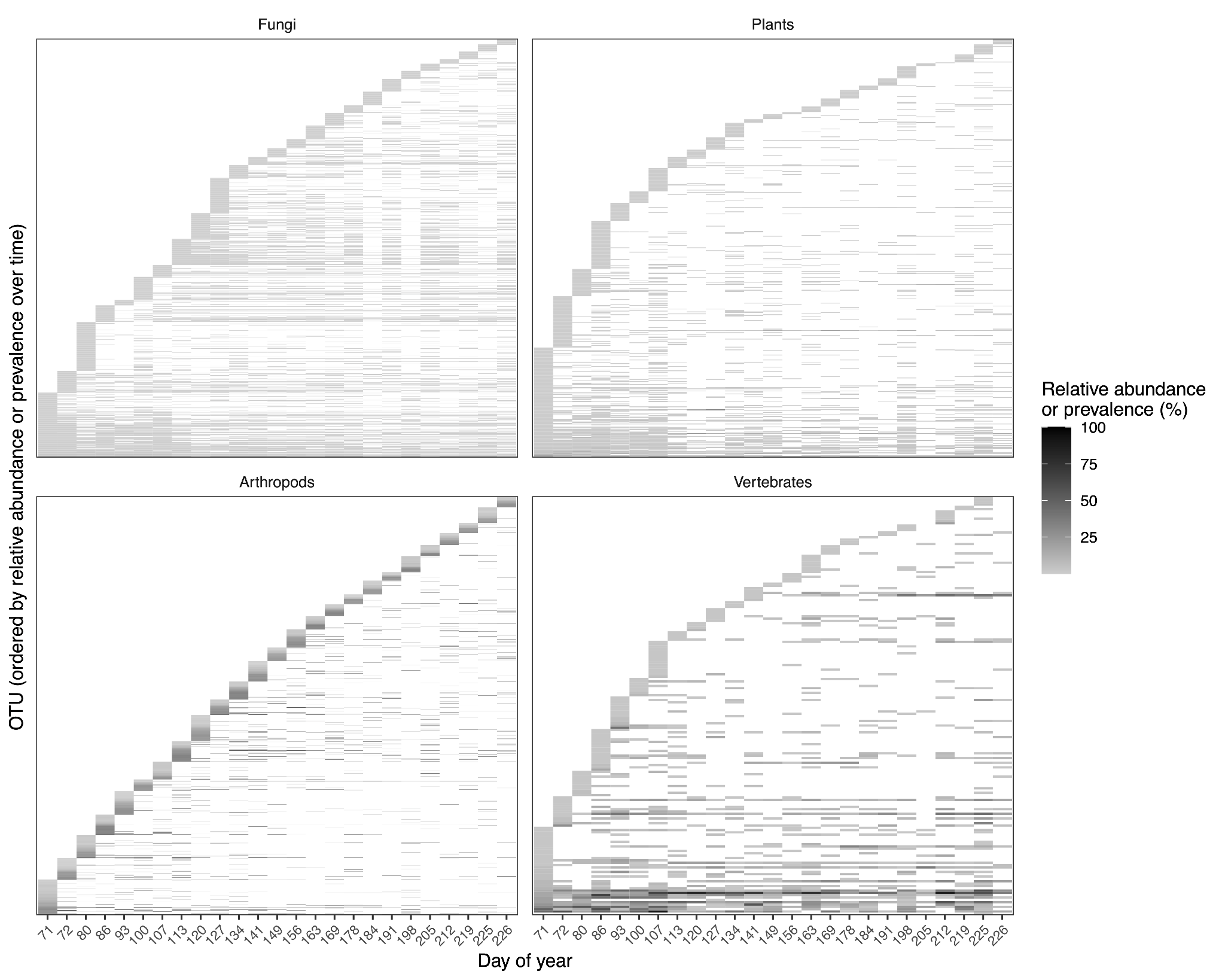

**Fig. S8.** Heatmap of OTU abundance (fungi and plants) and prevalence (arthropods and vertebrates) over time for each of the four primer sets. OTUs are ordered by the sampling period that they were first detected.

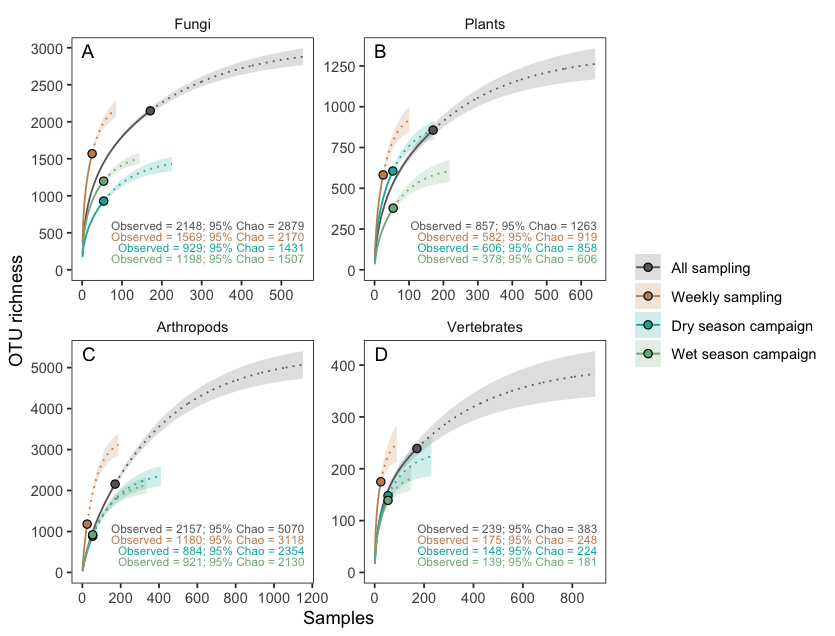

**Fig. S9.** OTU accumulation curves for (A) fungi, (B) plants, (C) arthropods, and (D) vertebrates across each of four BCI datasets: all sampling, weekly sampling, dry season campaign, and wet season campaign. Solid lines represent rarefaction from the observed richness (solid circles), and dotted lines represent extrapolation to 95% of the Chao1 index of asymptotic richness. Shading represents 95% confidence intervals.

Table S1. Comparison of number of OTUs between dry and wet season for each sample.

| Assemblage | Dry | Wet | Greater season | Greater pct | p |
| --- | --- | --- | --- | --- | --- |
| Fungi | 172.22 ± 4.82 | 215.43 ± 5.82 | Wet | 25.09 | 1.08E-07 |
| Plants | 73.79 ± 2.02 | 33.70 ± 1.88 | Dry | 118.94 | 9.36E-27 |
| Arthropods | 31.65 ± 1.44 | 30.48 ± 1.70 | Dry | 3.83 | 0.602 |
| Vertebrates | 17.52 ± 0.79 | 16.98 ± 0.97 | Dry | 3.16 | 0.667 |

Table S2. Comparison of number of OTUs between dry and wet season across all samples.

| Assemblage | Total | Dry | Wet | Shared | Shared pct | Greater season | Greater pct |
| --- | --- | --- | --- | --- | --- | --- | --- |
| Fungi | 1701 | 929 | 1198 | 426 | 25.04 | Wet | 28.96 |
| Plants | 729 | 606 | 378 | 255 | 34.98 | Dry | 60.32 |
| Arthropods | 1596 | 884 | 921 | 209 | 13.1 | Wet | 4.19 |
| Vertebrates | 196 | 148 | 139 | 91 | 46.43 | Dry | 6.47 |

Table S3. Comparison of airDNA concentrations (copies per extract) between wet and dry season for each assemblage.

| Assemblage | Dry | Wet | p |
| --- | --- | --- | --- |
| Fungi | 2.53e+10 ± 4.38e+09 | 2.99e+10 ± 2.27e+09 | <0.001 |
| Plants | 1.09e+06 ± 1.40e+05 | 2.05e+05 ± 2.89e+04 | <0.001 |
| Arthropods | 8.90e+08 ± 2.16e+08 | 5.94e+08 ± 1.06e+08 | 0.645 |
| Vertebrates | 4.31e+05 ± 9.35e+04 | 2.32e+05 ± 2.42e+04 | 0.07 |

Table S4. Primers used in this study.

| Gene | Forward primer name | Forward primer sequence (5'-3') | Reverse primer name | Reverse primer sequence (5'-3') | Amplicon size (bp) | PCR conditions |
| --- | --- | --- | --- | --- | --- | --- |
| ITS | ITSF1 | CTTGGTCATTTAGAGGAAGTAA | ITS2 | GCTGCGTTCTTCATCGATGC | 138-390 | Initial denaturation at 95 °C for 5 minutes, followed by 35 cycles of 45 seconds at 95 °C, 1 minute at 50 °C, and 90 seconds at 72 °C, and a final elongation at 72 °C for 10 minutes. |
| trnL | trnL-c | CGAAATCGGTAGACGCTACG | trnL-h | CCATTGAGTCTCTGCACCTATC | 110-201 | Initial denaturation at 94 °C for 3 minutes, followed by 40 cycles of 30 seconds at 94 °C, 30 seconds at 55 °C, and 1 minute at 72 °C, and a final elongation at 72° C for 10 minutes. |
| Short COI | ZBJ-ArtF1c | AGATATTGGAACWTTATATTTTATTTTTGG | ZBJ-ArtR2c | WACTAATCAATTWCCAAATCCTCC | 151-166 | Initial denaturation at 94 °C for 5 minutes, followed by 45 cycles of 30 s at 94 °C, 45 s at 45 °C, and 45 s at 72 °C, and a final elongation at 72 °C for 10 minutes. |
| Folmer COI | LCO1490 | GGTCAACAAATCATAAAGATATTGG | HCO2198 | TAAACTTCAGGGTGACCAAAAAATCA | 501-742 | Initial denaturation at 94°C for 3 minutes, followed by 35 cycles of 1 minute at 95°C, 1 minute at 40°C, and 75 seconds at 72°C, and a final elongation at 72°C for 7 minutes. |
| 12S | 12SVert_F | ACTGGGATTAGATACCCYACTATG | 12SVert_R | GAGRRTGACGGGCGGTDT | 375-407 | Initial denaturation at 94 °C for 3 minutes, followed by 45 cycles of 30 seconds at 94 °C, 30 seconds at 52 °C, and 1 minute at 72 °C, and a final elongation at 72 °C for 10 minutes. |
